# Fluorescence Lifetimes and Spectra of RPE and sub-RPE Deposits in Histology of Normal and AMD Eyes

**DOI:** 10.1101/2020.04.29.068080

**Authors:** Rowena Schultz, Kushmali C. L. K. Gamage, Jeffrey D. Messinger, Christine A. Curcio, Martin Hammer

**Affiliations:** Department of Ophthalmology, University Hospital Jena, Jena, Germany; Department of Ophthalmology and Visual Sciences, School of medicine, University of Alabama at Birmingham, Birmingham, Alabama, United States; Center for Medical Optics and Photonics, Univ. of Jena, Jena, Germany

**Keywords:** Age-related macular degeneration, retinal pigment epithelium, drusen, fundus autofluorescence, fluorescence lifetime, fluorescence spectra

## Abstract

**Purpose:** To investigate autofluorescence lifetimes as well as spectral characteristics of drusen and retinal pigment epithelium (RPE) in age-related macular degeneration (AMD).

**Method:** Fluorescence lifetimes and spectra of five eyes with AMD and nine control eyes were analyzed in cryosections by means of two-photon excited fluorescence at 960 nm. Spectra were detected at 490 – 647nm. Lifetime was measured using time-correlated single photon counting in two spectral channels: 500-550nm and 550-700nm. The fluorescence decays over time were approximated by a series of three exponential functions and the amplitude-weighted mean fluorescence lifetime τ_m_ was determined.

**Results:** 196 sub-RPE deposits were identified (AMD n=76, healthy n=120) and 230 RPE sites recorded. The peak emission wavelength of drusen was significantly green-shifted compared to RPE (peak at 570nm vs. 610nm), but not different between patients and controls. Drusen showed considerably longer τ_m_ than RPE: (ch1: 581 ± 163 ps vs. 177 ± 25 ps, ch2: 541 ± 125 ps vs. 285 ± 31 ps, p < 0.001). Drusen found in AMD eyes had longer lifetimes than drusen of controls (ch1: 650 ± 167 ps vs. 537 ± 145 ps, ch2: 600 ± 125 ps vs. 504 ± 111 ps, p < 0.001). In addition, drusen in AMD eyes showed a more homogenous fluorescence distribution and more drusen were larger than 63µm than in control eyes.

**Conclusions:** Ex vivo fluorescence imaging of drusen in cross-sections enables the separation of their autofluorescence from that of over- or underlying structures. Our analysis showed considerable variability of drusen lifetimes but not spectra. This indicates that drusen consist of a variety of different fluorophores or expose the same fluorophores to different microenvironments. Changes in drusen lifetimes could be related to AMD progression.

## Introduction

Although clinical fundus autofluorescence imaging (FAF) is a routine investigation technique in ophthalmology, especially in the diagnostics of age-related macular degeneration (AMD), the origin of the fluorescence signal is not fully understood yet. Different patterns of FAF are described and, in part, related to drusen and hyperpigmentation.^1, 2^ Recently, different fluorescence lifetime and spectral characteristics of drusen and retinal pigment epithelium (RPE) were revealed by fluorescence lifetime imaging ophthalmoscopy (FLIO).^3-5^ The latter in vivo investigation showed spectral differences between the fluorescence emission of drusen and surrounding retina/RPE. This difference may reflect different chemical composition: Whereas dominating fluorophores in the RPE are retinal-derived compounds,^6, 7^ drusen contain considerable fractions of lipids, phospholipids, and lipoproteins^8^. Fluorescence lifetime of drusen showed a wide variety, suggesting differences in their chemical composition.

As, however, FAF as well as FLIO provide en-face images with no depth resolution, fluorescence from various fundus layers is summed up in these signals. This overlay of diverse fluorophores from different anatomical structures makes it difficult to differentiate and characterize them. Thus, insight can be gained by complementing in vivo observations by histology. Marmorstein et al.^9^ investigated fluorescence emission spectra of RPE, sub-RPE deposits, and Bruch’s membrane upon excitation at different wavelengths. These authors reported shorter peak emission wavelengths for sub-RPE deposits than for lipofuscin in RPE cells. Also recent en face ex vivo imaging revealed different spectra for drusen and RPE.^10^ Measurements of histologic fluorescence lifetimes, as opposed to spectra, are limited to two eyes reported by Schweitzer et al., who found considerably longer lifetimes in drusen than in RPE.^11^

Therefore, in this study we measured fluorescence spectra and lifetimes on 196 drusen and 230 sites in RPE in order to check for differences between AMD and control eyes as well as small and larger drusen. This could give hints on fluorescence characteristics pointing to the development or progression of AMD. It would have been interesting to see whether macular and peripheral drusen show different fluorescence signatures. As, however, in our material only very few macular drusen were found, this question cannot be answered. As extramacular drusen are associated with age-related macular degeneration^12^ as well, the investigation of their fluorescence properties is considered meaningful too.^13-15^

## Methods

### Human donor tissue

Use of human tissues was approved by institutional review at UAB (#N170213002). The paired eyes of 9 Caucasian donors (mean age: 84.3 ± 3.4 years) were obtained (author CAC) from Advancing Sight Network (formerly the Alabama Eye Bank) for the purpose of independent studies that compared spectral and molecular characteristics of tissues processed fresh and -paraformaldehyde-fixed from the same donor.^16^ Before processing for histology, all eyes were opened anteriorly and subjected to an expert examination including post-mortem fundus inspection, ex vivo color fundus photography, and ex vivo optical coherence tomography (Spectralis, Heidelberg Engineering). From this post-mortem inspection, eyes were graded as unremarkable, AMD, or questionable AMD (small or infrequent deposits and RPE change). For this investigation, we distinguish only between AMD and controls, the latter comprising unremarkable eyes and eyes with questionable AMD.

To determine the effect of fixation on autofluorescence, two eyes from each donor were embedded together so that a tissue section from each eye was present on the same glass slide for subsequent analysis. From each donor eye, the cornea and a 2-mm-wide scleral rim was removed. For the left eye, the iris and lens were removed, the fundus photographed internally, then the tissue placed directly into carboxymethylcellulose (CMC, C9481 Sigma Aldrich, St. Louis, MO, USA) for freezing at -80°C. The right eye was preserved in 4% paraformaldehyde in 0.1M phosphate buffer for at least 24 hours before iris and lens removal, imaging, embedding in CMC next to the frozen left eye, and freezing at -80°C. Prior to freezing, the posterior poles of both eyes were trimmed to 14-mm-wide belts of retina, choroid, and sclera containing major landmarks (optic nerve head, fovea, and horizontal meridian of the visuotopic map) and extending anteriorly to pigmented tissue (ora serrata) at the edge of the ciliary body.

For diagnostic purposes, serial 10 µm cryosections were collected starting at the superior edge of the optic nerve head (of the preserved eyes). Sections were captured on pre-labeled 1×3 mm glass slides coated with 10% poly-L-lysine (Sigma Aldrich, St. Louis, MO, USA) and maintained at 37° during sectioning. To verify diagnosis and identify pathologic features of interest, every 20th slide was stained with periodic acid-Schiff-hematoxylin (K047 kit, Poly Scientific RD, Bayshore NY USA) to show basal laminar deposit, lipofuscin, and cell nuclei. Nearby unstained slides were shipped by overnight courier to Jena for fluorescence lifetime microscopy.

### Spectral and fluorescence lifetime imaging

Autofluorescence spectra and lifetimes were recorded using an inverted multiphoton laser scanning microscope (Axio Observer Z.1 and LSM 710 NLO, 63x oil immersion objective (Plan-Apochromat NA=1.4), Carl Zeiss, Jena, Germany) in combination with a femtosecond Ti:Saphire laser (Chameleon Ultra, Coherent Inc., Santa Clara, CA) and a single photon counting fluorescence lifetime imaging setup (Becker & Hickl GmbH, Berlin, Germany). The Ti:Saphire laser has a pulse repetition rate of 80 MHz with a pulse duration of 140 fs. The excitation wavelength for spectral and lifetime recordings was set to 960 nm.

The spectral QUASAR detector of the LSM710 has been utilized for spectral imaging. Backscattered excitation light was blocked by a beam splitter selecting the wavelength range 405 – 710 nm and the fluorescence emission was recorded in the range of 490 nm to 674 nm with a spectral resolution of 9.6 nm. All spectral images were recorded as an average of 40 fast scans (pixel dwell time of 1.6 µs) with a resolution of 512 x 512 pixel and a field of view of 96.4 x 96.4 µm.

The lifetime imaging measurements are based on the principle of time-correlated single photon counting (TCSPC). A single photon counting setup, consisting of two hybrid photomultiplier tubes (HPM-100-40) in non-descanned operation, each in combination with a SPC-150 TCSPC board, was used. Other components included are an optical beam splitter (LP555/BP500-550) to measure two-photon-excited fluorescence in two spectral channels (short spectral channel: SSC 500 – 550 nm and long spectral channel LSC: 550 – 700 nm). The excitation wavelength (960 nm) was cut off by a filter BG 39 (Schott, Mainz, Germany). All FLIM images have been acquired as an average of 90 fast scans (mean photon count per pixel of approximately 1000, pixel dwell time of 3.1 µs) with a resolution of 256 x 256 pixel and a field of view of 96.4 x 96.4 µm.

The fluorescence decay images from the FLIM detectors were analyzed using the software SPCImage 7.4 (Becker&Hickl GmbH, Berlin, Germany), which is described in detail elsewhere.^17^ The decays where approximated with a three-exponential model yielding in three decay time constants and three amplitudes. For further analysis, the intensity-weighted mean value of the time constants was calculated, denoted here as the mean fluorescence lifetimes (τ_m_).

For each image, regions of interests (ROIs = RPE or sub-RPE deposits/drusen) were selected in SPCImage and the average τ_m_ per ROI was used for further analysis. Drusen were categorized according to their size and appearance in fluorescence intensity images (graded by two independent investigators). Differences in homogeneity and shape were taken into account. Localization of the druse (macular or periphery) was not considered. Homogeneity was defined as homogenous drusen content with an even distribution of fluorescence with low variability. Non-homogenous drusen displayed many small internal patches of different fluorescence intensities. The drusen shape was either convex with a round silhouette and clear defined outlines to the overlaying RPE, or irregular with undefined contours and partially no clear separation to the RPE. Drusen diameter was measured as cross-sectional length along Bruch’s membrane. Statistical analysis was performed using SPSS 26 (IBM, SPSS Inc., Chicago, IL, USA). Distribution of drusen and size of drusen between control and healthy donor eyes was tested by the chi square test. Group mean values of lifetimes were compared by an unpaired t-test. Spectra were normalized to 1 by peak-normalization (division of all values by peak intensity value) for analysis. Differences of spectra between fixed and unfixed tissue, RPE and drusen, drusen size, as well as diagnosis were analyzed by fitting a quadratic polynomial mixed model with spectra and category (RPE vs. drusen, small vs. large drusen, or AMD vs. control) as fixed effects and random intercept. To estimate the differences, an interaction of spectra and category as well as spectra squared and category is included in the model.

## Results

### Drusen categories

Overall, 196 sub-RPE deposits/drusen were found in 16 of 18 donor eyes. Three RPE elevations were non-fluorescent and, thus, excluded from the analysis. Furthermore, the eyes of one donor did not show any signs of drusen and were excluded from further investigation. Five eyes were diagnosed with AMD by post mortem expert fundus inspection and OCT. Slightly more drusen were found over the whole section (macular and periphery) in AMD eyes: 76 drusen in five eyes (15.2 drusen per eye on average) vs. 120 drusen in 9 control eyes (13.3 drusen per eye). The diameter of the drusen showed a bi-modal distribution (fig. 1). Thus, in accordance with clinical diagnostic criteria, we distinguished between small drusen (N=140) with a diameter <63 µm and drusen larger than 63 µm in diameter (N=56). AMD eyes had more drusen >63 µm than controls (AMD: 34/76 drusen, 44.7% vs. control: 22/120 drusen, 18.3%; χ^2^ = 15.9, p < 0.000).

**Fig. 1:**
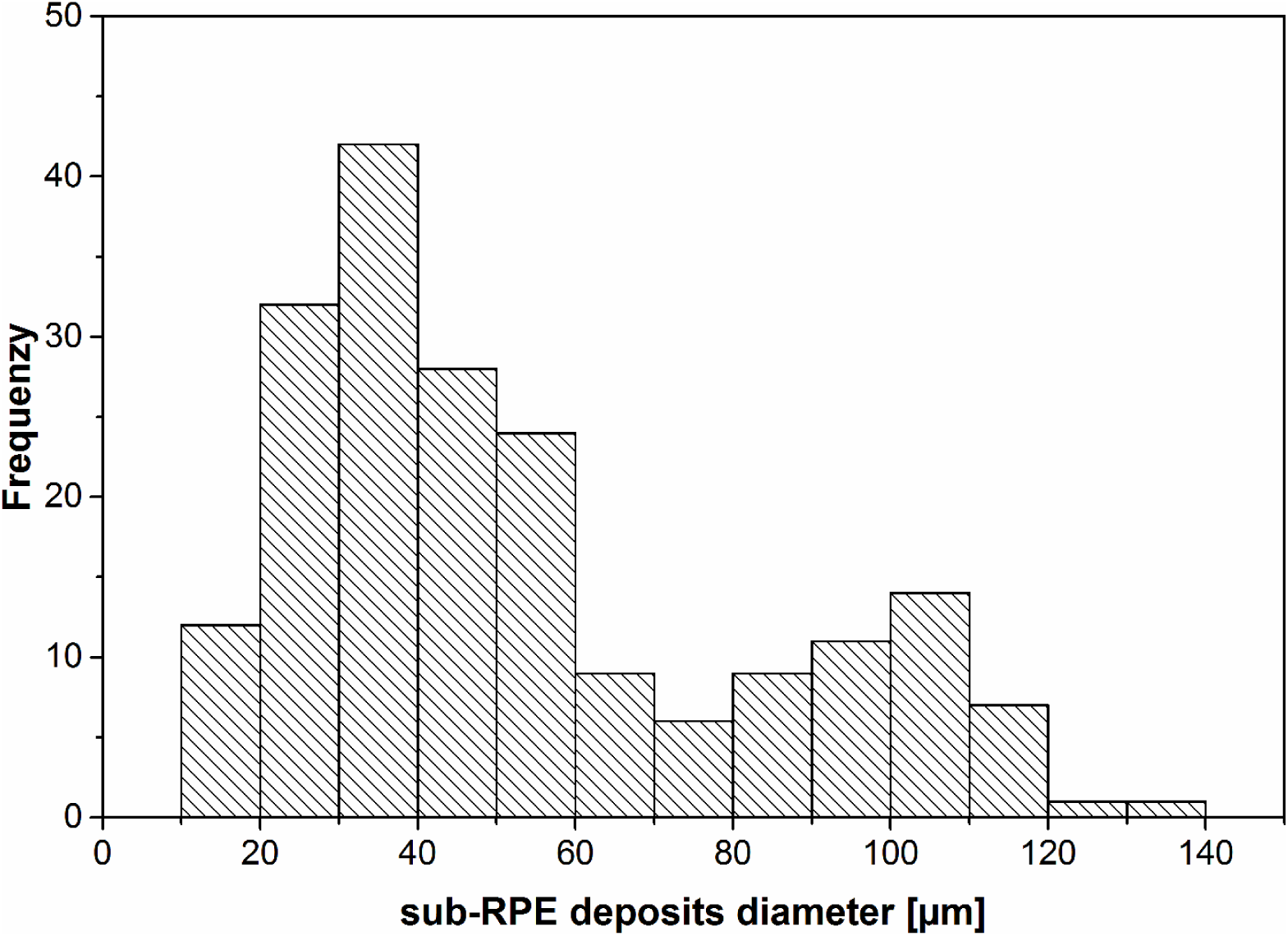
Histogram of sub-RPE deposits/drusen size.

We further distinguished the following categories of drusen according to their appearance in fluorescence intensity images: homogenous content (75/196 drusen, 38.3%, fig. 2A, B), non-homogenous content (121/196 drusen, 61.7%, fig. 2C, D), convex shape (86/196 drusen, 43.9%, fig. 2A,C), irregular shape (110/196 drusen, 56.1%, fig. 2B, D). Fig. 2 also shows fluorescence of Bruch’s membrane, which we did not quantify, since measurements on this small structure would have been greatly influenced by neighboring RPE, sub-RPE deposits and choriocapillaris.

**Fig. 2:**
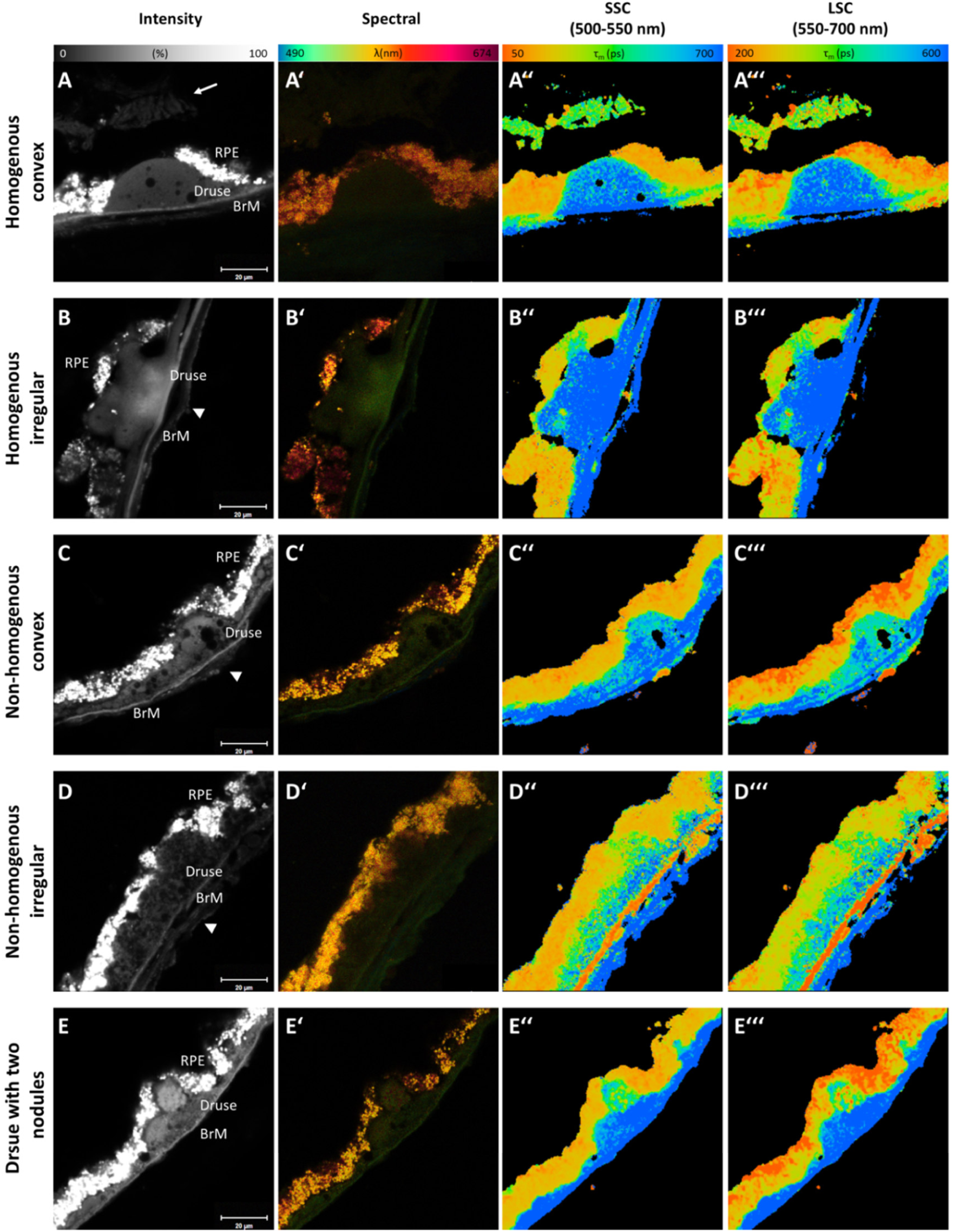
Examples of different sub-RPE deposits/drusen types, **A**) homogenous convex, **B**) non-homogenous irregular, **C**) Druse with two hyperfluorescent nodules (non-homogenous irregular), **D**) hyperfluorescent Druse (homogenous irregular). Intensity image, fluorescence emission spectra image and τ_m_ in SSC and LSC (pseudo-colored, SSC: 50-700 ps, LSC: 200-600 ps), Scale bar = 20 µm, arrow = retina (disrupted preparation artifact), arrowhead = choroid (disrupted preparation artifact).

### Fluorescence lifetimes

First we checked for the influence of tissue fixation on the fluorescence lifetimes. As these did not significantly differ either for RPE or for drusen (table 1), we did not distinguish between fresh frozen samples and those which were PFA-fixed before deep freezing and slicing.

**Table 1:** Mean fluorescence lifetimes τ_m_ for RPE and drusen in fresh frozen and fixed tissue. Mean ± standard deviation, fresh frozen samples N = 111 for RPE and 98 for drusen, fixed samples N = 119 for RPE and 98 for drusen, results for the short-(SSC) and long-wavelength spectral channel (LSC) are given, statistic by unpaired Students t-test.

|  | fresh frozen | fixed | p |
| --- | --- | --- | --- |
| RPE SSC | 175 ± 22 | 178 ± 28 | 0.288 |
| RPE LCS | 288 ± 27 | 283 ± 35 | 0.238 |
| Drusen SSC | 572 ± 140 | 589 ± 184 | 0.46 |
| Drusen LSC | 540 ± 110 | 542 ± 139 | 0.91 |

Drusen showed longer lifetimes (SSC: 581 ± 163 ps, LSC: 541 ± 125 ps, N = 196) than RPE (SSC: 177 ± 25 ps, LSC: 285 ± 31 ps, N = 230). These differences were highly significant (p < 0.001) for both channels. Whereas RPE fluorescence lifetimes were relatively homogeneous, drusen lifetimes showed a high variability (fig. 3). The lifetimes of drusen as well as that of the RPE were significantly longer in the AMD eyes compared to drusen and RPE in control eyes (table 2).

**Table 2:** Mean fluorescence lifetimes τ_m_ for RPE and drusen in control and AMD donor eyes. Mean ± standard deviation, control N = 146 for RPE and 120 for drusen, AMD N = 84 for RPE and 76 for drusen, results for the short-(SSC) and long-wavelength spectral channel (LSC) are given, statistic by unpaired Students t-test.

|  | control | AMD | p |
| --- | --- | --- | --- |
| RPE SSC | 167 ± 22 | 193 ± 21 | <0.001 |
| RPE LCS | 274 ± 26 | 304 ± 31 | <0.001 |
| Drusen SSC | 537 ± 145 | 650 ± 167 | <0.001 |
| Drusen LSC | 504 ± 111 | 600 ± 125 | <0.001 |

**Fig. 3:**
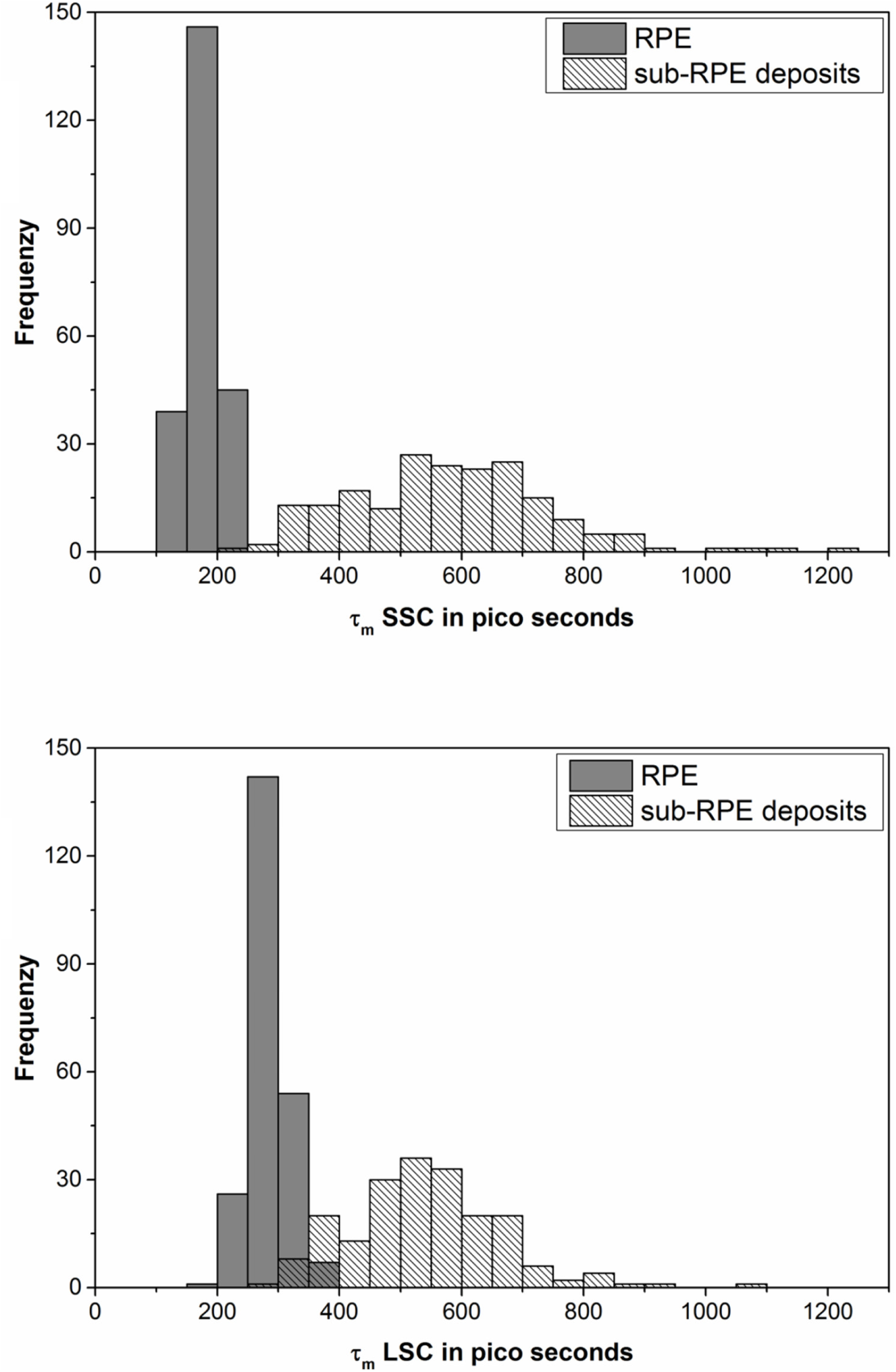
Histograms of τ_m_ for RPE and sub-RPE deposits/drusen, top: SSC, bottom LSC.

Homogeneous drusen are more abundant in AMD patients (AMD: 56/76 drusen, 73.7%, control: 19/120 drusen, 15.8%). They had longer lifetimes compared to non-homogenous drusen in general (SSC: 657 ± 174 ps vs. 534 ± 138 ps, LSC: 605 ± 136 ps vs. 502 ± 100 ps, p < 0.001 for both). Especially in AMD patients homogenous drusen had significantly longer lifetimes compared to the non-homogenous ones. This, however, did not hold for the control subjects (results of subgroup comparisons are listed in table 3). The shape of the drusen was not differently distributed in AMD and healthy subjects and had no influence on the lifetimes (data not shown).

**Table 3:** Mean fluorescence lifetimes τ_m_ for homogenous and non-homogenous drusen (according to their appearance in intensity images). Mean ± standard deviation, control N = 19 for homogenous and 101 for non-homogenous drusen, AMD N=56 for homogenous and 20 for non-homogenous drusen, results for the short-(SSC) and long-wavelength spectral channel (LSC) are given, statistic by unpaired Students t-test.

|  | control | AMD | p |
| --- | --- | --- | --- |
| Homogenous Drusen SSC | 555 ± 160 | 691 ± 165 | 0.003 |
| Homogenous Drusen LSC | 533 ± 145 | 629 ± 125 | 0.007 |
| Non-homogenous Drusen SSC | 533 ± 143 | 535 ± 111 | 0.959 |
| Non-homogenous Drusen LSC | 499 ± 103 | 517 ± 78 | 0.473 |

As mentioned above, we differentiated two size ranges of drusen according to clinical diagnostics. Larger drusen had significantly longer lifetimes than the small ones (SSC: 662 ± 171 ps vs. 548 ± 149 ps, LSC: 600 ± 128 ps vs. 518 ± 116 ps, p < 0.001 for both).

### Fluorescence emission spectra

Averaged spectra are shown in fig. 4. No difference was found between the spectra of fixed and fresh-frozen tissue (not shown). Therefore, spectra from both preparations were pooled. The fluorescence emission of drusen occurred at shorter wavelength than that of RPE. As the RPE fluorescence peaks at 610 nm, the peak of the drusen fluorescence was found at 570 nm (fig 4 A). This difference of RPE and drusen spectra was significant. We found an interaction of category (drusen or RPE) and spectra (p = 0.032) as well as category and spectra squared (p = 0.017) in the polynomial mixed regression model. The RPE spectra were not significantly different for AMD donor eyes and controls, although a slight shift of the rising edge (between 500 and 600 nm) towards shorter wavelengths was observed for AMD donors (fig. 4 C). The same holds for the drusen spectra. Although fig. 4 D shows a slight hypsochromic shift for the AMD donors, the peak was at 570 nm for both groups and the spectra were not significantly different. Similarly, the comparison of drusen >63 µm with smaller ones showed a slight hypsochromic shift of the larger drusen but no significant difference (fig. 4 B).

**Fig. 4:**
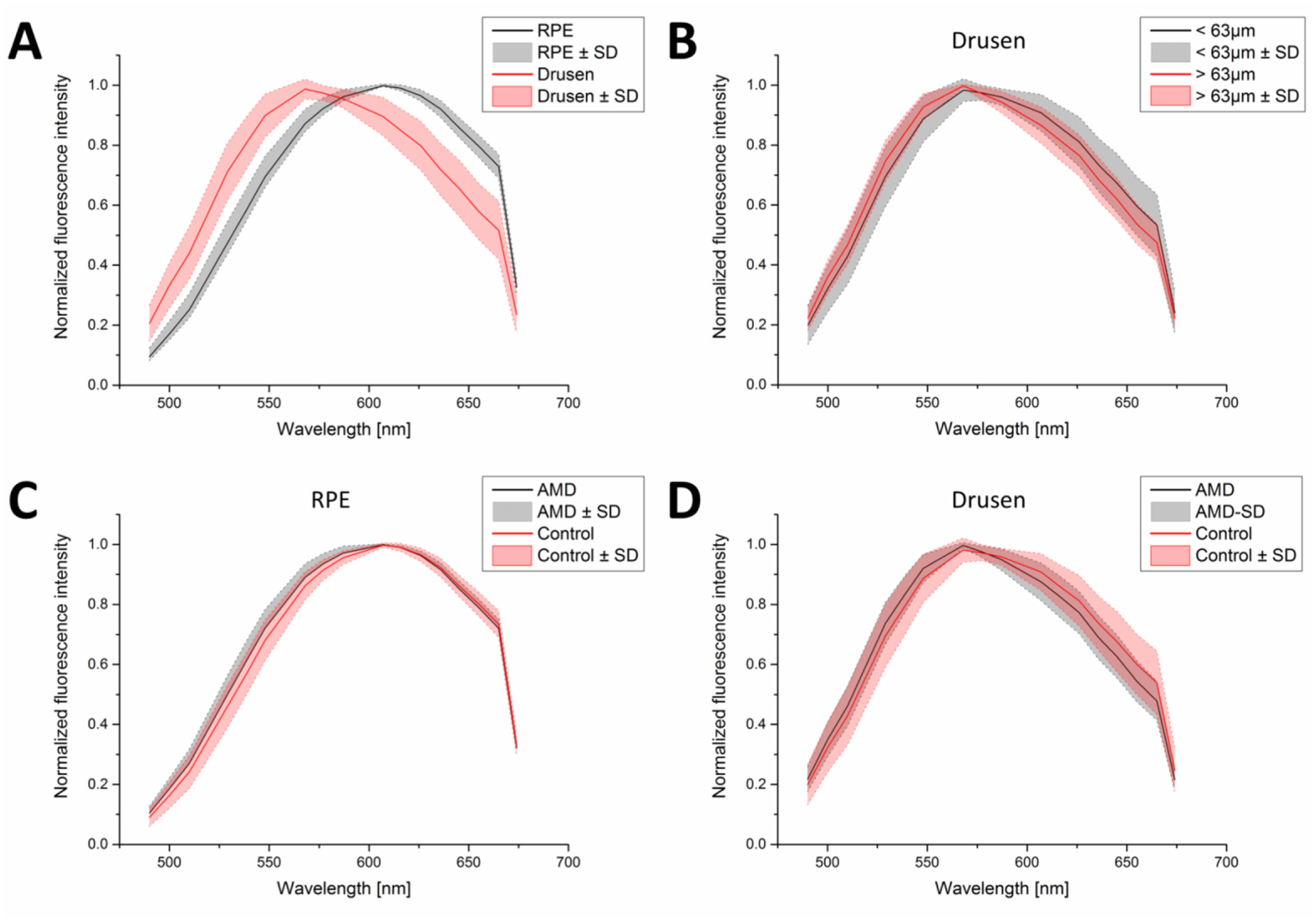
Fluorescence emission spectra averaged over groups of measurements, filled areas indicate the range of standard deviation. **A**: spectra of all drusen (red) and RPE (black), **B**: drusen <63 µm (black) and >63 µm (red) in diameter, **C**: RPE of AMD patients (black) and controls (red), **D**: drusen of AMD patients (red) and controls (blue).

## Discussion

Clinical FAF, excited at 488nm, is thought to originate from lipofuscin and melanolipofuscin in RPE cells.^18-20^ In our investigation, as in prior studies,^9^ we found histologic fluorescence also from Bruch’s membrane and sub-RPE deposits such as basal laminar deposits, basal linear deposits (BlinD) and drusen, however, the RPE fluorescence was by far the strongest. Furthermore, RPE fluorescence clearly showed the granular structure, known from previous investigations with two-photon^21, 22^ as well as super-resolution microscopy,^23^ which points to lipofuscin and melanolipofuscin granules. Their fluorescence was spectrally similar to that reported for lipofuscin and melanolipofuscin.^10, 24^ We found the peak of the RPE spectrum at 610 nm, as was reported by Delori et al. for human post mortem RPE excited at 510 nm.^18^ In vivo, these authors found peak emissions at 631 nm and 621 nm upon 510 and 470 nm excitation, respectively. Thus, slight differences in the emission peak might be due to the excitation conditions (wavelength and 960 nm two vs. one-photon excitation). A dependence of the peak emission on the excitation wavelength was also found by Marmorstein et al. in histology.^9^ These authors measured slightly shorter emission peaks for 488 nm excitation (approximately corresponding to our 960 nm two-photon excitation), which might be due to the fact that they used exclusively macular samples. Also, Delori reported shorter emission peaks for the macula than measured at an eccentricity of 7° upon 470 nm excitation in vivo.^18^ On the other hand, BenAmi et al. did not find differences in the spectral signature of foveal, perifovea, and near-periphery histologic RPE samples when decomposing fluorescence spectra using non-negative matrix factorization (NMF).^24^ But also the preparation protocol may have an influence and two-photon excitation may act somewhat different than one photon excitation.

For Drusen in vivo, Arend et al. found a hypsochromic spectral shift in areas with drusen in vivo.^25^ They revealed a spectrum peaking at 560 nm, which they attributed to drusen fluorescence. We measured very similar drusen fluorescence with a peak at 570 nm. Tong et al. reported a drusen-specific spectrum with a peak most abundantly found at 520 nm (excitation 480 nm).^10^ This is shorter than what we measured. However, Tong et al. derived their drusen spectrum from measured spectra by NMF; here we present peak-normalized spectra which were directly extracted from the measurements. Furthermore, we used histologic cross sections whereas Tong et al. applied en-face imaging, which could yield in fluorescence contribution also of Bruch’s membrane, which is known to have short-wavelength fluorescence.^24^ Taken together, this might explain the difference of drusen spectra found in both investigations.

Our fluorescence maximum of drusen was 40 nm shorter than that of RPE, which is in agreement with our recent in vivo findings using the two spectral channels of FLIO^5^ and indicates that drusen contain other fluorophores than lipofuscin. Although lipofuscin-like inclusions in drusen may occur occasionally, their number is very small^26-28^ and should not have a measurable impact on the total drusen fluorescence. We found only slightly hypsochromic shifted spectra of RPE and drusen in AMD donors, compared to controls, as well as of drusen larger than 63 µm in diameter vs. smaller ones upon unchanged peak emission wavelengths. Thus, the diagnostic relevance of these spectral differences remains to be determined.

Fluorescence lifetime, however, seems to have high potential for diagnosis. The fluorescence lifetime of the RPE in our study was significantly longer in AMD donor eyes in both spectral channels. This was in agreement with the finding of prolonged fluorescence lifetimes in AMD patients in vivo.^4^ The in vivo lifetimes, reported in that paper, are to some extent longer in general compared to our ex vivo findings presented here. This might have several reasons. First, although our measurements were close to the in vivo data, we cannot exclude postmortem changes or alterations by the preparation procedure. Second, the in vivo data, recorded from en face images, certainly contain fluorescence components from other tissues than RPE, which might have longer lifetimes. Third, fluorescence excitation by different lasers (femtosecond laser for two-photon excitation in our measurements, one-photon excitation with a picosecond laser in vivo), might make a difference.

Drusen are the major intraocular risk factor for progression to advanced AMD (neovascularization and atrophy)^15, 29, 30^ and may be predictive for the disease progression.^21-23^ Traditionally, drusen were identified and classified by fundus inspection or color fundus photography, respectively, and AMD staging greatly relies on the number, size, and morphology of drusen.^31^ More detailed insight in different types of drusen localization and morphology is gained by OCT.^32-38^ Several studies investigated the autofluorescence of fundus areas with drusen. Whereas Delori et al. found hypo-fluorescence of the drusen center, often surrounded by a hyper-fluorescent annulus,^39^ Göbel et al. found drusen with increased, decreased, as well as unremarkable autofluorescence intensity.^40^ Investigating refractile drusen, Suzuki et al. found a transition from uniform hyper-FAF to a ring of hyper-FAF, presumably due to a phase of photoreceptor shortening followed by loss of the RPE on the top of the druse.^41^ Accordingly, FLIO has revealed a wide variety in drusen fluorescence lifetimes.^3, 4^ In a recent study, one third of soft drusen was found to be hyperfluorescent. Over all drusen, no difference of lifetimes compared to adjacent retinal areas was found.^5^ This could be due to multiple stages of druse evolution.^41, 42^

In in vivo FAF and FLIO imaging, contributions to the fluorescence signal from all retinal layers have to be considered. Accordingly, Delori et al.^39^ explained the central hypofluorescence of drusen with a hyperfluorescent annulus by the thinning of the RPE on top of the druse with a translocation to its rim. Longer fluorescence lifetimes in some of the drusen^4, 5^ and a general shift towards shorter emission wavelengths compared to the adjacent fundus^5^, however, indicate an independent autofluorescence contribution from the drusen themselves. Although the drusen fluorescence, in general, is weaker than that of the RPE, its contribution to the total FAF in in vivo en face imaging can be considerable, if the drusen are much thicker than the RPE which is a cellular mono-layer.

In order to distinguish fluorescence emissions from different fundus layers, in this study we investigated histological cross sections. This clearly revealed different fluorescence properties of RPE and drusen. Although drusen can be virtually non-fluorescent, most of them emit fluorescence at shorter wavelength and with longer decay times than the RPE. Thus far, this study confirms in vivo FLIO results and an earlier investigation performing spectrally resolved FLIM in the histology of one AMD and one control donor eye^11^. Relative to our previous study, ^11^ in histologic cross-sections druse fluorescence is not overlaid by that of the RPE, and we had more donor material. First of all, we grouped drusen by their size for comparison of fluorescence parameters. According to clinical AMD classification criteria,^30, 31^ we distinguished drusen <63µm which were clearly localized and round, from larger more widespread sub-RPE deposits. The latter, more extended drusen showed significantly longer fluorescence lifetimes, indicating a different fluorophore composition. This difference was clearly underlined by finding some drusen containing two hyperfluorescent nodules (fig. 2E) with clearly distinct lifetimes. On the other hand, longer lifetimes in the larger drusen seem to contradict our in vivo findings, where we observed no difference in large (soft) drusen compared to their surrounding fundus^5^. However, it has to be kept in mind that this holds for hyperfluorescent drusen in vivo only. Also here we found highly fluorescent drusen (fig. 2B) having long lifetimes. Furthermore, a comparison with the in vivo fluorescence is difficult for small drusen (<63 µm) as these are on the resolution limit of clinical FLIO.

Besides difference in the shape of the drusen (round or stretched at Bruch’s membrane, in our study not associated with AMD) and all heterogeneity in fluorescence lifetime, drusen lifetime was significantly longer in donors with AMD than in aged donors lacking clear AMD pathology. Furthermore, the abundance of drusen, especially homogenous ones, was higher in the AMD eyes and the percentage of drusen larger than 63 µm was higher than in the controls. In OCT, the most common pattern observed for drusen in eyes of AMD patients was a convex shape with a homogeneous filling.^43^ In histology, drusen also significantly differ with respect to their ultrastructural morphology and clear differentiation between homogenous structures and highly heterogeneous ones are possible.^28, 44^ Soft drusen are described to be made of homogeneous content ^28^ and have been reported to highly correlate with AMD and AMD progression.^8^ One could speculate that the homogenous drusen with long lifetimes, found in AMD patients, are soft drusen. Our findings underscore that drusen are linked to AMD pathology. However, the higher lifetime of drusen in AMD also indicates that a different drusen composition might hint to the disease.

Drusen and BlinD are extracellular accumulations of lipids, proteins, and minerals. Plasma LDL and HDL deliver essential lipophilic compounds such as vitamins A and E, lutein, and cholesterol to the RPE and outer retina.^45-47^ Furthermore, apoA, B, and E lipoproteins, secreted from the RPE, may bind to Bruch’s membrane.^48, 49^ Subsequently, those lipoproteins may degrade and fuse to a lipid pool, according to Curcio et al. analogue to an oil spill.^8, 50^ Unfortunately, little is known about the fluorescence of such lipids upon blue light or 2-photon IR excitation respectively. As, however, drusen formation shares some similarity with that of atherosclerotic plaques, where blood LDL binds to the extracellular matrix of the vessel wall^51, 52^, a comparison with fluorescence investigations in those plaques is interesting.^53^ Park et al. found an emission at 550nm from the lipids in the plaques, which was distinguishable from the fluorescence of collagen and elastin^54^. This lipid fluorescence showed a long lifetime of 5.3ns. This lifetime is considerably longer than the one we measured for drusen. That could be due to different instrumentation and fluorescence excitation conditions (Park et al. used a Nd:YAG laser, 355nm/1ns). On the other hand, drusen might be more complex as atherosclerotic plaques as not only serum derived lipids contribute to their formation, but also molecules from photoreceptor degradation as well as visual pigment turnover forming lipoprotein-derived debris.^8, 50^

Several limitations of this study have to be mentioned: There was no clinical data or diagnosis available for the donor eyes. Thus, classification in AMD or control had to rely on post-mortem fundus inspection and OCT. Furthermore, only small and intermediate drusen were found in the samples. However, it cannot be excluded that larger soft drusen were present initially, but were lost in the tissue preparation and slicing procedure as soft drusen are biomechanically fragile. Also it was not always possible to separate drusen and BlinD fluorescence from that of basal laminar deposits.

The measured drusen size might have been underestimated due to off-axis cuts that missed the largest part of the deposit. As however, despite this systematic underestimation of the drusen size, a significant difference of fluorescence lifetimes for drusen, measured smaller or larger than 63 µm, was found, this has to be considered as a matter of fact. This study used 2-photon-excitation of the fluorescence because this gives a better spatial resolution, especially of the FLIM images recorded by detectors at the non-descanned port of the microscope. This, however, limits somewhat the comparability with in vivo FLIO measurements which use 1-photon picosecond fluorescence excitation. Nevertheless, in this study we found similar lifetimes as in vivo. Although fixation, in agreement with Delori et al.^18^, seems to have only minor impact on fluorescence spectra as well as lifetimes, it has to be pointed out that post-mortem changes of fluorophore composition and properties of the embedding matrix may have occurred.

Additionally, preparation of the donor eyes for cryosectioning may alter fluorescence lifetimes by a change of the viscosity of the sample while they are frozen. This all limits the comparability of our results to in vivo findings. However, the lifetimes, found for RPE, are similar to that reported from in vivo studies. Thus, we assume that also the prolongation as well as the diversity of drusen fluorescence lifetimes reflects the in vivo situation at least qualitatively.

In conclusion, we found considerably longer fluorescence lifetimes for drusen than for RPE. Furthermore, the lifetime was longer for larger drusen and drusen with homogenous fluorescence patterns. The lifetime of drusen as well as RPE was longer in AMD eyes than in controls. Drusen emitted fluorescence at shorter wavelengths than RPE. If, however, fluorescence spectra are different for AMD patients and controls remains questionable.

